# Offspring performance is well buffered against stress experienced by ancestors

**DOI:** 10.1101/2020.01.10.901421

**Authors:** Yifan Pei, Wolfgang Forstmeier, Bart Kempenaers

**Affiliations:** Department of Behavioural Ecology and Evolutionary Genetics, Max Planck Institute for Ornithology, 82319 Seewiesen, Germany

**Keywords:** early developmental stress, trans-generational effect, anticipatory effect, condition transfer, resilience, morphology, reproductive performance, lifespan, quantitative genetics, multiple testing

## Abstract

Evolution should render individuals resistant to stress and particularly to stress experienced by ancestors. However, many studies report negative effects of stress experienced by one generation on the performance of subsequent generations. To assess the strength of such transgenerational effects we used a strategy aimed at overcoming the problem of type I errors when testing multiple proxies of stress in multiple ancestors against multiple offspring performance traits, and applied it to a large observational data set on captive zebra finches (*Taeniopygia guttata*). We combined clear one-tailed hypotheses with steps of validation, meta-analytic summary of mean effect sizes, and independent confirmatory testing. With this approach we assess to what extent offspring performance in adulthood depends on (1) direct effects of own experiences during early development, (2) indirect condition-transfer effects of the early environment experienced by the parents and the grandparents, and (3) beneficial effects of a match between the environments experienced by the offspring and by its parents. Our study shows that drastic differences in early growth conditions (nestling body mass 8 days after hatching varied 7-fold between 1.7 and 12.4 gram) had only moderate direct effects on adult morphology (95%CI: r=0.19-0.27) and small direct effects on fitness traits (r=0.02-0.12). In contrast, we found no indirect effects of parental or grandparental condition (r=-0.017-0.002; meta-analytic summary of 138 effect sizes), and mixed evidence for small benefits of matching environments, as the latter was not robust to confirmatory testing in independent data sets. This study shows that evolution has led to a remarkable robustness of zebra finches against undernourishment and that transgenerational effects are absent.

**Author Summary:** How the early life conditions of your ancestors might influence your own life, an aspect of epigenetic inheritance, has become a popular topic among evolutionary biologists and has sparked much interest by the general public. Many theoretical and empirical studies have addressed this question, leading to theories of adaptive programming and condition transfer and ideas of epigenetic or genetic organization. Despite the popularity of this topic, however, there is a lack of a standard framework to guide empirical studies, which are at risk of over-interpreting the most significant effects that might emerge by chance alone when conducting a large number of tests. In this study, we used long-term observational data on multiple morphological and life-history traits of hundreds of male and female zebra finches with information on the early life conditions of both the focal birds and their parents and grandparents. This allows us to comprehensively quantify the magnitude of direct and transgenerational effects of early developmental conditions. Our study (1) proposes a standardized statistical framework for future investigations, (2) summarizes the average effect size (in zebra finches) and indicates the sample sizes needed to pick up such an effect, and (3) provides a counter statement to a growing faith in the ubiquity of transgenerational effects despite their limited evolutionary or mechanistic plausibility.

## Introduction

Early developmental stress experienced by an individual may have long-term negative effects on its morphology, physiology, behavior and reproductive performance later in life (i.e. ‘direct effect’) [1–7]. Effects of conditions in early life could also be transmitted to subsequent generations [8–12], potentially via the inheritance of epigenetic markers (e.g. altered DNA methylation, transmission of small interference RNAs or hormones) [13–15]. Such inheritance of acquired traits could exist either because of an inevitable transfer of condition from one generation to the next (i.e. ‘condition transfer’ or ‘carry-over’ or ‘silver-spoon’ effect, e.g. low-condition mothers produce low-condition offspring) [11,16–19], or because organisms have evolved mechanisms of adaptive programming, where offspring were ‘primed’ by their parents and perform the best if they grow up in an environment similar to that of their parents (i.e. ‘anticipatory effect’, hypothesis of matching / mismatching environments) [10,20,21]. It is important to distinguish between the two types of trans-generational effects, i.e. (1) ‘condition transfer’ and (2) ‘anticipatory effects’, in experiments (e.g. match/mismatch), especially when the unavoidable intra-generational (3) ‘direct effects’ of early conditions experienced by the individual itself are substantial [10–12,17].

From an evolutionary perspective, we would expect that natural selection acts to minimize the susceptibility of organisms to harmful direct and indirect, condition-transfer effects. Fitness-related traits in particular are selected to be well-buffered against detrimental influences from the environment (evolution of stress tolerance, robustness and developmental canalization; e.g. [22, 23]) and selection will disfavor mothers that handicap their own offspring. In general, detrimental carry-over effects may be inevitable to some extent, but selection will work against them. In contrast, ‘transgenerational anticipatory effects’ are thought to have evolved because they have an adaptive function. Such “transgenerational anticipatory programming” of offspring may have evolved when the environments in which parents and offspring grow up are generally similar, e.g. [20, 21], and when proximate mechanisms of epigenetic inheritance enable it, e.g. [13–15]. Studies of epigenetic inheritance boomed since the early 1990s, focusing mostly on organisms such as fungi and plants that are immobile and lack the differentiation between soma and germ cells [14, 24], but also on nematodes, fruit flies, mice and humans [15,25–34]. However, in the latter group of organisms, the evolution of proximate mechanisms that would allow an adaptive programming seems less plausible, given that the environment experienced by the soma would need to be conveyed to the germ line [14]. In sum, the widespread existence of both types of transgenerational effects seems somewhat unlikely, because condition transfer is selected against and anticipatory effects may lack a plausible mechanism.

Although the mechanisms behind most of the observed epigenetic inheritance remain largely unclear [14, 35], evolutionary biologists have studied transgenerational effects and have estimated the fitness consequence of stress experienced by one generation on individuals of subsequent generations in various animal systems [36, 37], sometimes with individuals from the wild [38], but mostly with captive-bred animals e.g. [21,39–42]. Such effects have typically been investigated experimentally across two generations, i.e. effects of increasing stress experienced by the parents on the offspring, using brood or litter-size manipulation [39, 40], restricted food supply during female pregnancy or nestling or puppy stages [43], restraint stress exposure during early life where individuals were intermittently deprived from social interactions [44], corticosterone intake during female pregnancy or early individual development [45], and cold or heat shock (mostly for insects, e.g. in *Drosophila* and *Tribolium* [46, 47]). In general, the reported significant effects are often accompanied by numerous non-significant test results, and sometimes involve a flexible interpretation of the direction of the effect. Moreover, transgenerational effects were sometimes reported as sex-specific (interaction effect between the sex of the parent and that of the offspring). For example, in humans, effects from (grand-) mother to (grand-) daughters and from (grand-) father to (grand-) sons have been reported [48, 49]. To assess the importance of these effects, we suggest that rigorous testing of *a priori* hypotheses and meta-analytic summary of effect sizes are needed.

Here, we use observational data of >2000 captive zebra finches from a long-term, error-free pedigree to study the sex-specific effects of multiple stressors experienced during early development on later-life morphology and fitness-related traits. We consider both direct, intra-generational effects and effects of developmental stress experienced by parental and grandparental generations. For all individuals, we systematically recorded variables that have previously been used as indicators of early developmental conditions (brood size [2,39,50], hatching order [42,51,52], laying order of eggs [53, 54] and clutches [55], egg volume [56], and nestling body mass at 8 days old [3]), as well as morphological traits (tarsus [2, 39]and wing length [21,39,42], body mass [2,21,42], abdominal fat deposition[3], beak color[2,3,42,57]) as dependent variables. For a subset of birds, we also measured lifespan (N = 821 individuals) and aspects of reproductive performance (female clutch size in cages (N = 166 females) and in aviaries (N = 274 females), female fecundity (N = 230 females), male infertility in cages (N = 132 males) and in aviaries (N = 237 males), male siring success (N = 281 males), female embryo mortality (N = 228 genetic mothers), nestling mortality for a given social mother (N = 233) and for a given social father (N = 228), female and male seasonal recruits (N = 126 males and N =125 females, details see Methods), e.g. [2,21,42,45,58].

Regarding condition transfer, we focus our analyses on the *a priori* hypothesis that the stress that an individual’s parents and grandparents experienced in early life has detrimental effects on the morphology and reproductive performance of that individual as an adult. We assume that the direction of effects is independent of the sex of the focal individual. We further hypothesize *a priori* that if such transgenerational effects were sex-specific, the environment experienced by mothers and grandmothers would affect daughters and grand-daughters whereas the environment experienced by fathers and grandfathers would affect sons and grand-sons. Such one-tailed expectations have the advantage that trends which are opposite to the expectation can be quantified as negative effect sizes. If the null hypothesis is true, i.e. if there is no effect, we expect a meta-analytic mean effect size that does not differ from zero. Regarding anticipatory effects, we focus on the *a priori*, one-tailed hypothesis that offspring perform better as adults when they experienced similar early-life conditions as their parents.

First, we validate the six proxies of early developmental stress by examining their direct effects on the individual itself. Second, we use meta-analysis to average transgenerational effect sizes across multiple traits reflecting either morphology or reproductive performance of adult male and female zebra finches. Lastly, we use an independent dataset to assess whether the significant findings from the initial tests can be replicated.

## Results

We examined the effects of six variables describing early developmental conditions (potential stressors) on ten measures of morphology and on 13 aspects of reproductive performance, resulting in 138 predictor-outcome combinations (6 x 23 tests). We thus obtained 138 effect sizes for the direct effects (intra-generational; Table S1), 828 effect sizes for the inter-generational condition-transfer effects (i.e. effects of the early-life experiences of the six ancestors: two parents and four grandparents, 6 x 138; Table S1) and 46 effect sizes for the anticipatory effects (i.e. effects of similarity in nestling mass between mother and offspring and between father and offspring for 23 performance traits; Table S2).

### Validation of stressors using direct effects

Of the six putative indicators of early developmental conditions only one measure had significant consequences for the adult individual (Fig 1). Nestling body mass measured at 8 days of age affected both adult morphology (mean r=0.229, 95%CI: 0.186 to 0.272, P<0.0001) and reproductive performance (mean r=0.070, 95%CI: 0.021 to 0.119, P<0.0001; Fig 1, Table S3). The mass of nestlings when 8 days old varied by a factor of 10 (range: 1.2-12.4 g, N=3,525 nestlings), and light-weight nestlings had a clearly reduced chance of survival to adulthood (see Fig S1). Among the survivors (N = 3,326) and among those individuals included in the analyses of direct and transgenerational effects (N = 2,099), mass at day 8 still varied by a factor of 7 (range: 1.7-12.4 g).

**Fig 1.**
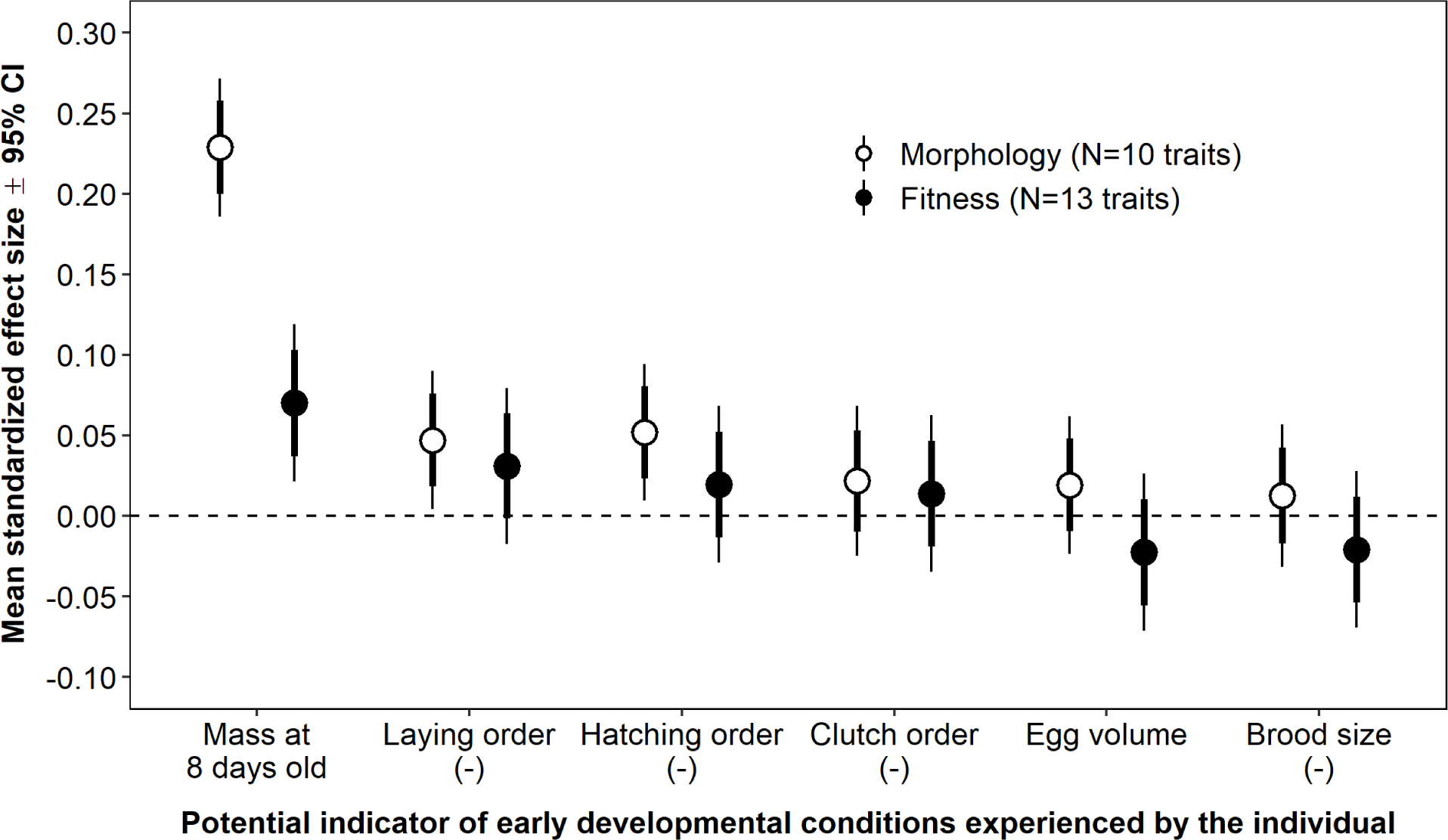
Average magnitude of direct effects of six potential indicators of early developmental conditions experienced by the individual itself on their adult morphology (averaged across 10 traits; open symbols) and fitness-related traits (13 traits; filled; Table S3). Error bars show two types of 95% CIs: thick lines refer to the single estimate and thin lines are Bonferroni adjusted for conducting 12 tests (figure-wide significance). Morphological traits (sex specific body mass, tarsus length, wing length, fat score and beak colour, median N = 944 females and 1008 males) were measured when individuals were 93-229 days old. Fitness-related traits include male and female lifespan, male and female number of seasonal recruits, female clutch size (in cages and aviaries) and fecundity, male fertility (in cages and aviaries), male siring success in aviaries, female embryo survival, and female and male nestling survival (median N = 228 individuals). Four out of the six indicators of early-life conditions were multiplied by −1 (indicated by (-)) such that positive effect sizes reflect better performance under supposedly better conditions. Morphological and fitness-related traits as well as indicators of early-life conditions were Z-transformed to yield effect sizes in the form of Pearson correlation coefficients.

Other indicators of developmental conditions, despite being widely used as proxies in the published literature, had little direct effect on the individual later in life. Therefore, in the following analyses we only use nestling body mass at day 8 as the proxy of early-life condition of parents and grandparents to assess the strength of the two types of transgenerational effects.

### Transgenerational effects of nestling mass: condition transfer

We did not find any evidence for a transgenerational effect of nestling mass either of the parents or of the grandparents on the adult offspring (mean estimate of 138 transgenerational effects after accounting for some level of non-independence between the response variables r=-0.007, 95% CI: −0.017 to 0.002; Table S4; see also Fig 2B and 2D; Table S5). Among the many correlations examined, only one was significant: the nestling mass of the mother correlated positively with the reproductive performance (clutch size in cages and aviaries, fecundity in aviaries, embryo survival, nestling survival, seasonal recruits and lifespan; Table S1) of her daughters (mean r=0.071, 95% CI: 0.025 to 0.117, P=0.003, without correction for multiple testing, Table S5; also see Fig 2D). This finding was mostly driven by a large positive effect of maternal early growth on daughter fecundity (Fig 3F), which was even larger than the direct effect of the daughters’ own nestling mass at day 8 (Fig 3E). For all other dependent traits that were influenced by nestling mass, the direct effects (Fig 3A and 3C) exceeded the indirect maternal effects (Fig 3B and 3D).

**Fig 2.**
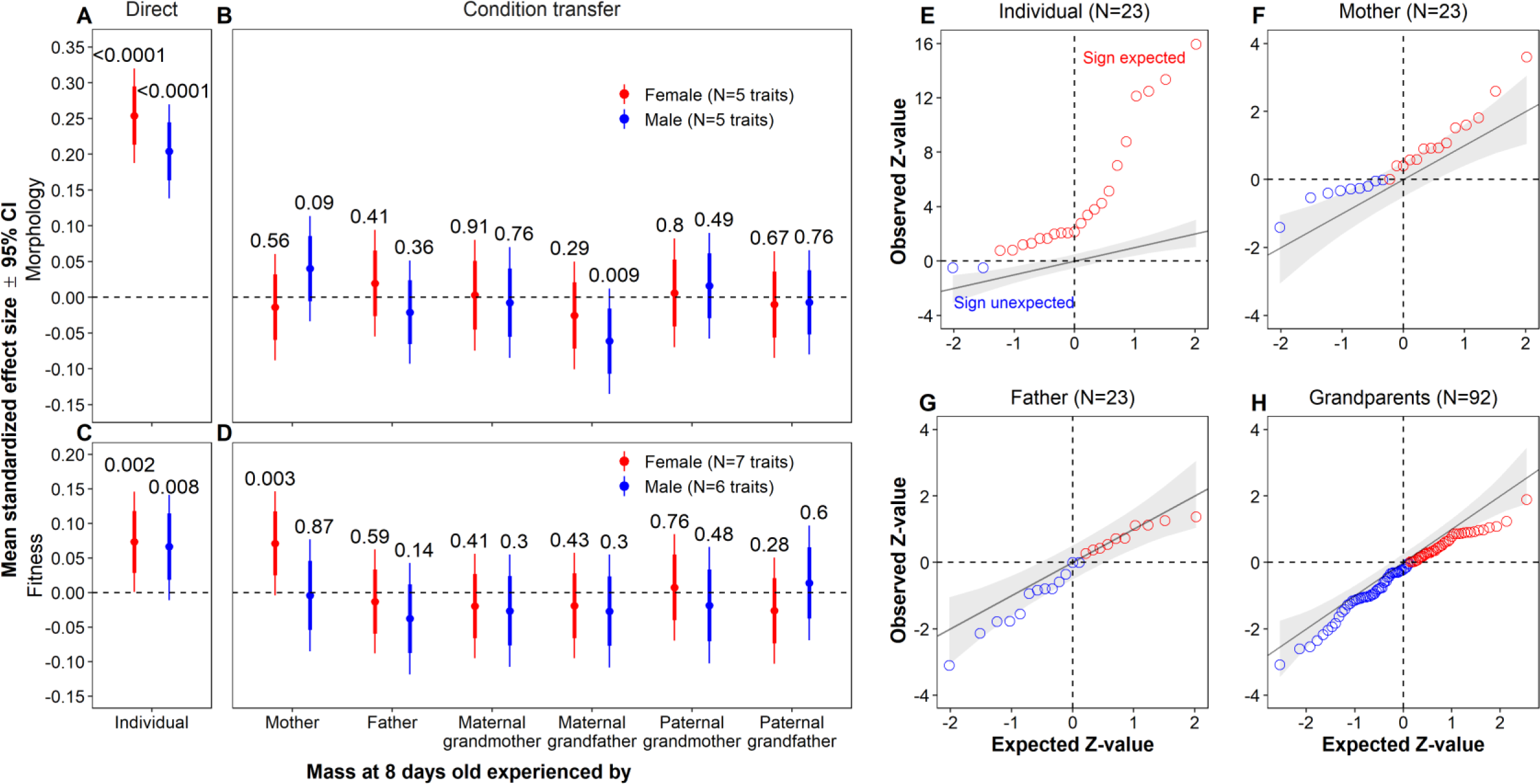
Trans-generational condition transfer effects of early developmental conditions (measured as nestling body mass at 8 days of age). (A-D) Average magnitude of condition transfer effects from 6 types of ancestors (B, D) in comparison to the direct effects of the experience of the individual itself (A, C) on morphological (mean of 5 traits; A, B) and fitness-related traits (mean of 6 or 7 traits; C, D) for individual females (red) and males (blue; Table S5). Error bars show two types of 95% CIs: thick lines refer to the single estimate and thin lines are Bonferroni adjusted for conducting 28 tests (figure-wide significance among A-D). Indicated P-values refer to each average effect estimate without correction for multiple testing. For further explanations see legend of Fig 1. (E-H) ZZ-plots of expected versus observed Z-values of the effects of early developmental conditions (mass at 8 days old) experienced by the focal individual itself (E), its mother (F), its father (G) and its four grandparents (H) on 10 morphological and 13 fitness-related traits. N indicates the number of tests. Red indicates that the sign of the estimate is in the expected direction, blue indicates that the sign is in the opposite direction. Lines of identity (where observation equals prediction) and their 95% CIs are shown.

**Fig 3.**
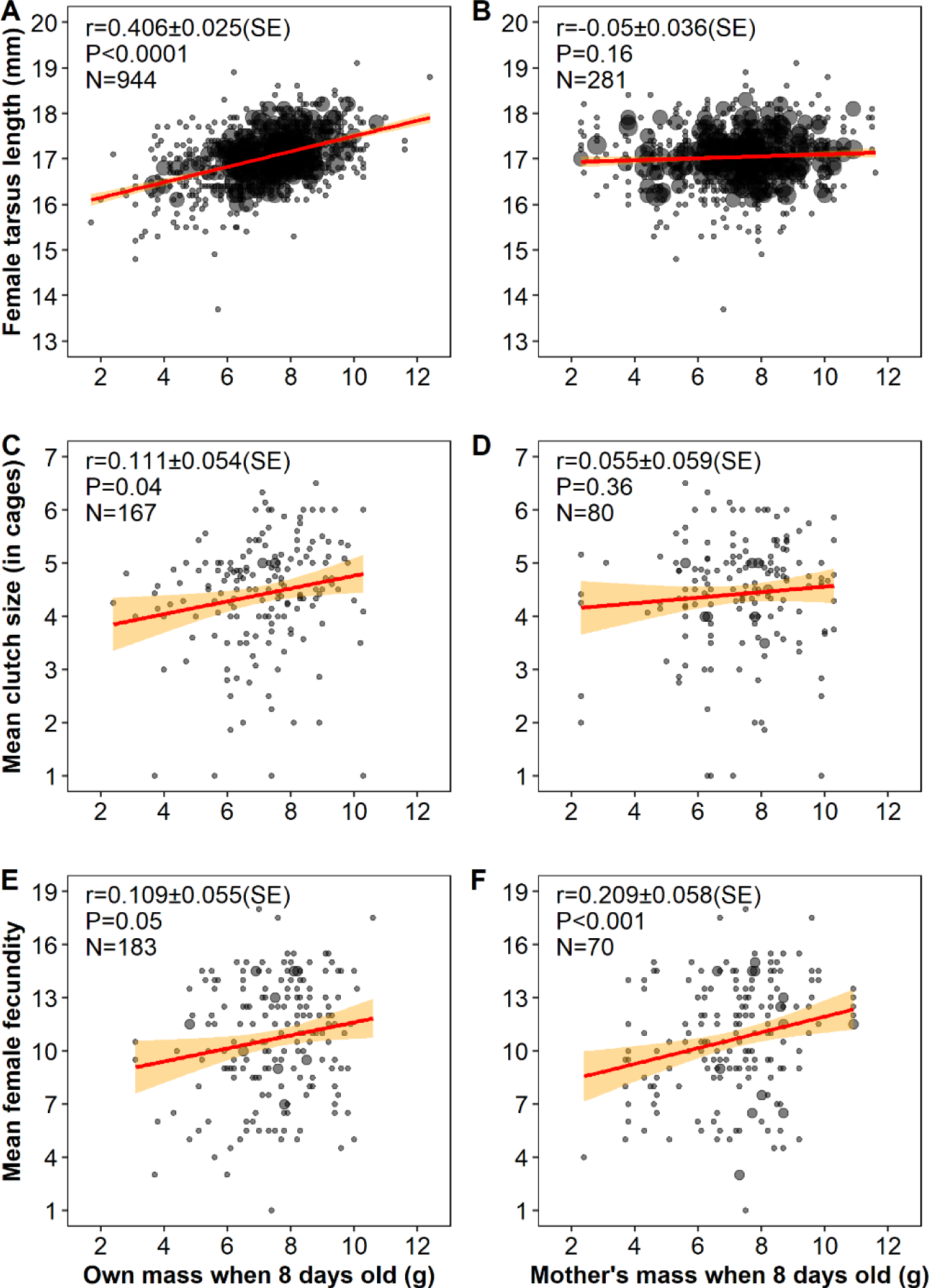
Comparison of direct effects (left column) and effects of condition transfer from mother to daughter (right column). Relationship between nestling mass at 8 days old experienced by the focal female (A, C, E) or by her mother (B, D, F) and tarsus length (A, B), mean clutch size in cage breeding (C, D) and mean female fecundity when breeding in an aviary (E, F). Dot size reflects sample size. Red lines are the linear regression lines (± 95% CI, orange shading) on the data shown, while indicated effect sizes (r with SE, P, N: number of females for A, C, E, and number of mothers for B, C, D) reflect estimates from mixed-models. Note that each individual’s own mass when 8 days old corresponds to one value of the dependent variable whereas each mother’s mass at 8 days old can correspond to multiple values of the dependent variable (one for each of her daughters).

The direct effects of nestling mass on the individual’s adult traits are clearly stronger than expected under a random distribution of effect sizes (Fig 2E; see also Fig 2A and 2C). In contrast, the positive effects of the early-life condition of the mother (Fig 2F) are not much stronger than the presumably coincidental negative effects (opposite to expectations) of the early-life condition of the father and the grandparents (Fig 2G and 2H; see also Fig S2). The two significant maternal effects (upper right corner in Fig 2F) are those on daughter fecundity (see above, Fig 3F) and on daughter clutch size (r=0.103, 95% CI: 0.025 to 0.181, P=0.01 without correction for multiple testing, Table S1). These findings are not independent, because clutch size and fecundity are strongly correlated (r=0.71, N=230 females, Fig S3 and Table S6), partially due to the fact that they are measured in the same breeding season (N=183 females).

### Transgenerational effects of nestling mass: anticipatory effects

Offspring performed significantly better when growing up under similar conditions as their parents (similarity in mass at day 8), but the effect size was small (mean estimate of 46 transgenerational effects after accounting for some level of non-independence between the response variables r=0.028, 95%CI: 0.017-0.039; Table S7; see also Fig 4 and Table S8). This was mainly driven by the positive effects of (1) father-daughter similarity on the daughters’ size (i.e. tarsus and wing length) and (2) mother-daughter similarity on the daughters’ fitness-related traits (Fig 4 and Table S8; see also Table S2).

**Fig 4.**
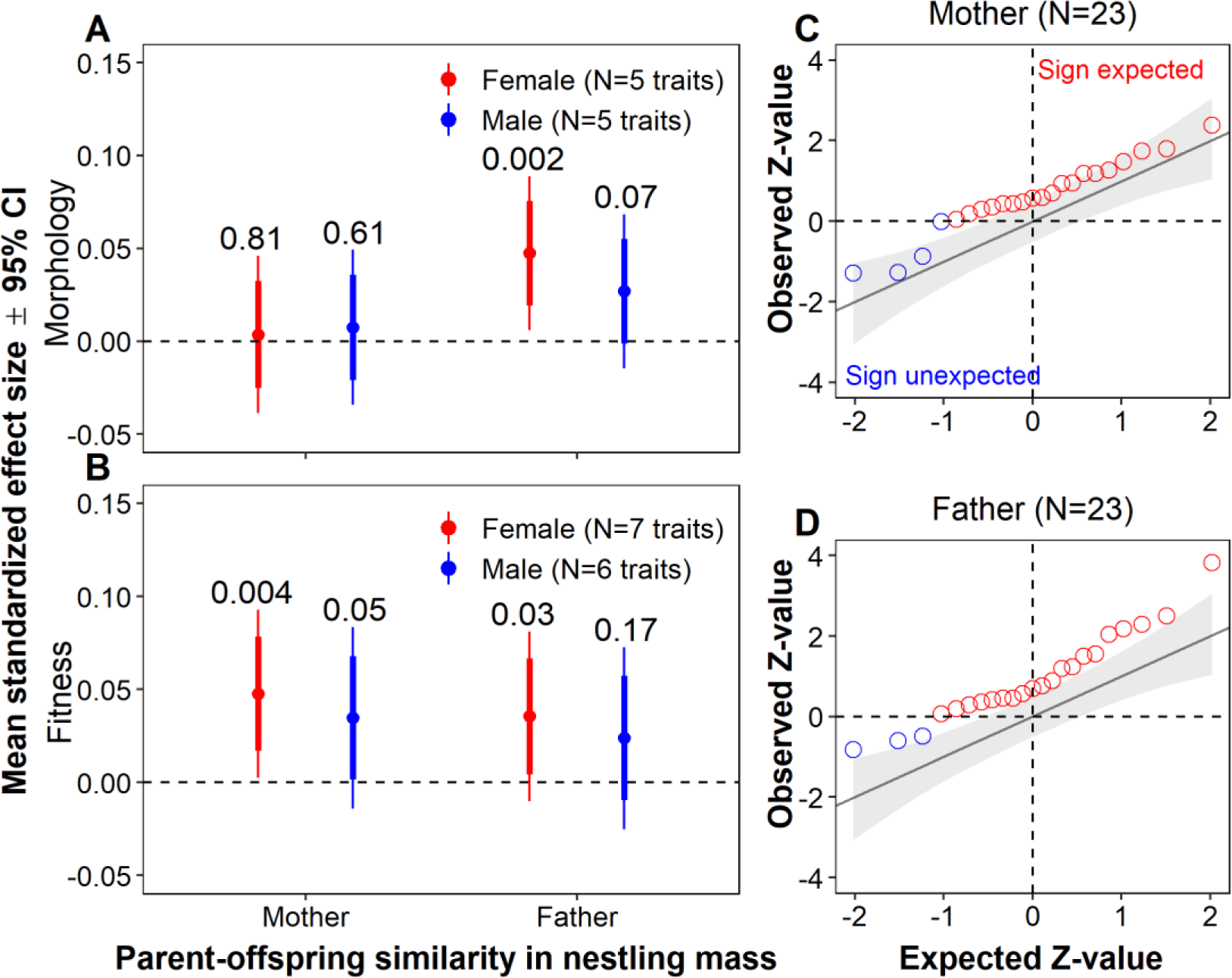
Transgenerational anticipatory effects (of similarity in early growth conditions between parents and offspring). (A-B) Average magnitude of anticipatory effect on morphological (mean of 5 traits; A) and fitness-related traits (mean of 6 or 7 traits; B) for individual females (red) and males (blue; Table S8). Error bars show two types of 95% CIs: thick lines refer to the single estimate and thin lines are Bonferroni adjusted for conducting 8 tests (figure-wide significance in A-B). Indicated P-values refer to each average effect estimate without correction for multiple testing. (C-D) ZZ-plots of expected versus observed Z-values of the effects of similarity in conditions (mass at 8 days old) between the focal individual itself and its mother (C), and between the focal individual itself and its father (D) on 10 morphological and 13 fitness-related traits (Table S2). For further explanations of A-B see legends of Fig 1 and C-D see legends of Fig 2.

### Confirmatory tests on independent data

To independently verify the strongest and most plausible findings of (1) condition transfer from the mother affecting daughter fecundity, (2) anticipatory effects of similarity between mother and daughter in their nestling body mass on daughter fecundity and (3) anticipatory effects from the father-daughter similarity on daughter body size, we examined an independent data set. This confirmatory data set consists of females with incomplete data on transgenerational effects – missing data on nestling mass of grandparents –not used in the initial tests, population ‘Seewiesen’) and of birds from two additional populations with shorter pedigrees (i.e. a recently wild-derived population ‘Bielefeld’ and a domesticated population generated from interbreeding between populations ‘Krakow’ and ‘Seewiesen’, population ‘Krakow’, see Methods for details). From this dataset, we used all available measures of female fecundity (including the three strongly correlated traits fecundity in aviaries, clutch size in cages and clutch size in aviaries) and size (including the two strongly correlated traits tarsus and wing length; see Table S6, Fig S3). All effect sizes of the confirmatory analysis are listed in Table S9. For all three tests, the initial effect size was clearly larger than the independent verification effect size (exploratory versus confirmatory, details in Fig 5 and Table S9) and besides the effect of father-daughter similarity on daughter tarsus length none of the confirmatory tests was significant.

**Fig 5.**
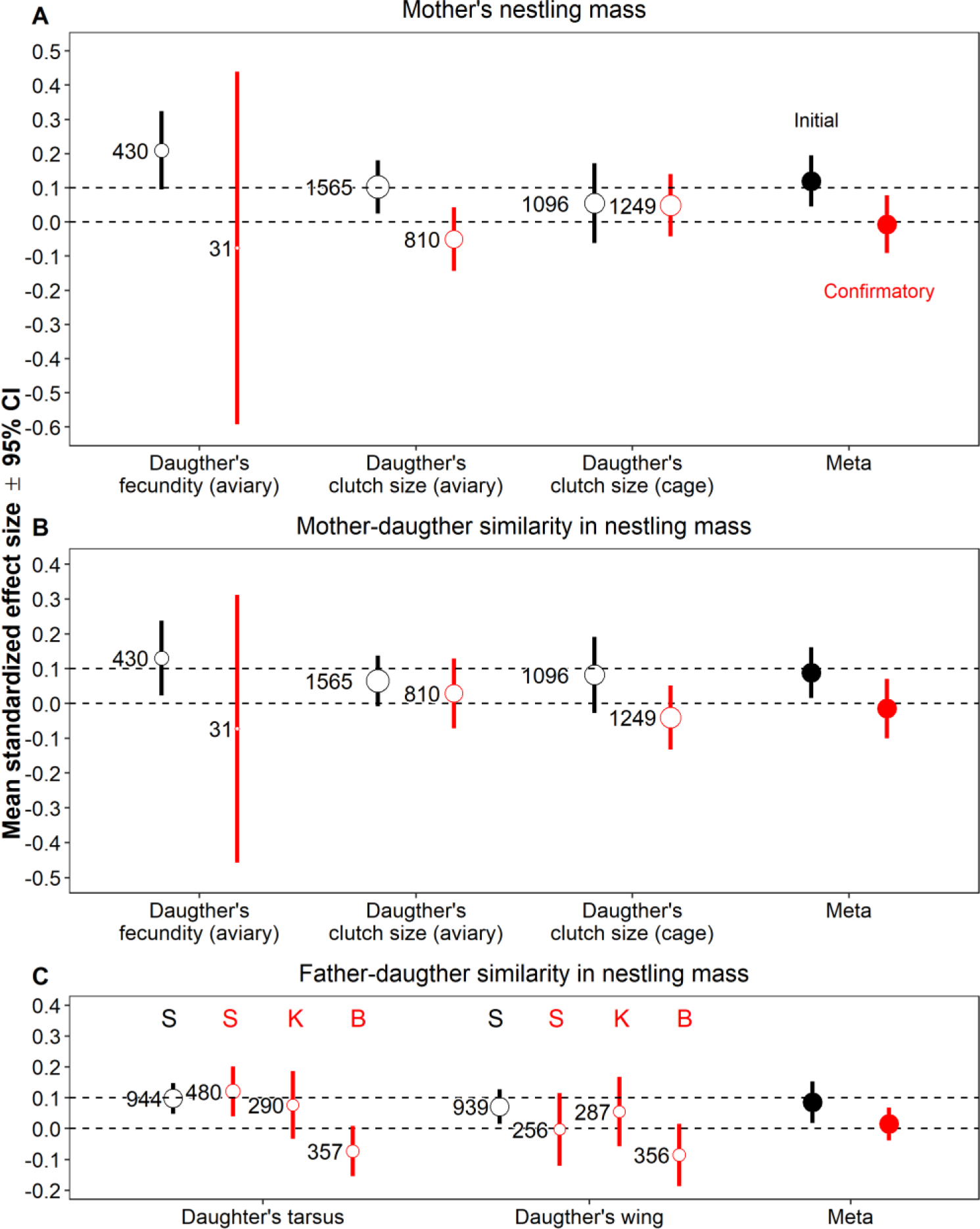
Confirmatory analysis of transgenerational condition transfer (A) and anticipatory (B-C) effects. Effect sizes (mean ± 95% CI without controlling for multiple testing) of mother’s mass at 8 days old (A) and the similarity between mother and daughter in their mass at 8 days old (B) on daughters’ fecundity, clutch size in cage and aviary (open) and the similarity of father and daughter in their mass at day 8 on the daughters’ tarsus and wing length (open symbols; C). The filled symbols show the meta-summarized effect sizes from the initial data set (black) and from the confirmatory data set (red). Numbers in the plots refer to the number of individuals (tarsus and wing length), clutches (clutch size in aviary and cage) or breeding seasons (fecundity). Daughter fecundity-related and size-related traits, mother’s mass at 8 days old, and similarity between mother-daughter and father-daughter in nestling mass were Z-transformed to yield effect sizes in the form of Pearson correlation coefficients. Tarsus and wing length were analysed by population due to the between-population difference in body size (C), where ‘S’, ‘K’ and ‘B’ refer to populations ‘Seewiesen’, Krakow’ and ‘Bielefeld’. Additional details see Table S9.

## Discussion

Our study supports the general idea that individuals are resilient to stress and particularly to stress experienced by ancestors. Even though individuals differed 7-fold in body mass when 8 days old, nestling mass only had small effects on morphology and reproductive success later in life. Our results clearly reject the hypothesis of condition transfer between generations, in line with the idea that selection acts against transmitting a handicap to the next generation. We found some evidence for transgenerational anticipatory effects, but the mean effect was small (r = 0.028), and did not hold up in an independent confirmatory test (Fig 5B and 5C). These mixed results indicate that the effect size for transgenerational anticipatory effects must be exceedingly small [59, 60]. In conclusion, transgenerational effects were absent or miniscule and direct effects on fitness traits were relatively small given that some of the offspring were seriously undernourished. Thus, the notion of organismal robustness seems more noteworthy than the claim of sensitivity to early-life conditions within and across generations, which dominates the literature (e.g. [2,8–10,21,42]). This begs the question whether the underrepresentation of studies emphasizing “robustness” in the literature is the result of the predominating framework of hypothesis testing, where the rejection of the null hypothesis is almost a pre-condition of getting published [61].

We found that direct intragenerational effects of early environment on morphology were of moderate magnitude while effects on fitness-related traits were small, which is largely in line with previous findings [2, 7]. Regarding transgenerational effects of early stress, we examined the existing zebra finch literature [21,39,40,42,45,58] and found that studies typically report a large number of tests (median number of discussed combinations of stressors, traits and sex: 18, range: 7-150). Only 15% of all tests were statistically significant, which is not far from the random expectation, especially if some non-significant findings were not reported. Additionally, an experimental study on zebra finches found no transgenerational anticipatory effect [21] and a meta-analysis of studies on plants and animals found no effect of transgenerational condition transfer [59]. A more recent meta-analysis found stronger transgenerational effects [62], yet this may be a consequence of selective inclusion of large effects. In three out of three randomly checked studies included in this meta-analysis [45,63,64], we found that only 38%, 75% and 50% of reported effects had been included, respectively.

Given the small (expected) effect sizes, we argue that transgenerational effects can sensibly only be studied within a framework that ensures a comprehensive reporting of all effect sizes and a meta-analytic summary of these effects. Focus on a subset of tests (e.g. those that are significant) leads to bias, but selective attention may be advisable in two situations. Firstly, when there is an independent selection criterion. For example, we limited our analysis of transgenerational effects to those involving only the most powerful indicator of early developmental conditions. In this case, the selection criterion (magnitude of direct effects, Fig 1) was established independently of the outcome variable (magnitude of transgenerational effects). Secondly, when there is an independent data set. For example, we selected the largest transgenerational effects from a first data set, and assessed them independently using a second data set (Fig 5). Consistent with the phenomenon of the winner’s curse [65], we found that selective attention to large effects yields inflated effect size estimates compared to the independent replication.

Selective attention to large effects makes the published effect size estimates unreliable. Thus, we propose to base conclusions on meta-analytic averages of all effect sizes that have been judged worth of investigation before any results were obtained. With this approach we shift our attention from identifying the supposedly best predictor and best response towards the quantification of the magnitude of an average predictor on an average response. Clearly, the latter is more reliable than the former, just as the average of many numbers is more robust than the maximum. Accordingly, the meta-analytic summary yields narrow confidence intervals around the estimated mean effect size. Note, however, that the estimated 95% CI might be somewhat anti-conservative (i.e. too narrow), because the summarized effect sizes are not fully independent of each other (multiple response variables are correlated; see Fig S3 and Table S6). In the cases where we summarize a large number of effect sizes (138 estimates in Table S4 and 44 estimates in Table S7) we fitted a random effect that controls for some of this non-independence, and this led to confidence intervals that are about 20% wider (compared to dropping the random effect). This approach of modelling and quantifying the degree of non-independence cannot be applied when summarizing only few effect size estimates (between 2 and 13 estimates in Figs 1, 2, 4, and 5), meaning that the indicated confidence intervals will be somewhat too narrow.

In our study, five out of six putative indicators of early developmental stress had little or no direct effect on an individual’s morphology and fitness later in life (Fig 1). This suggests that it is not worth to examine these traits for transgenerational effects ([14]; see also Fig S2), unless one can plausibly assume that some indicators of early stress cause direct effects, while others cause transgenerational effects. This differs from previous studies that showed various direct effects (e.g. of brood size [2,66,67], laying order [53, 68] and hatching order [42]), but did not attempt to meta-summarize all examined effects. In contrast to the other five variables, nestling mass (8 days old) was clearly associated with both nestling survival (Fig S1) and adult performance. However, its strongest effects were on morphology (highest r=0.41, Fig 3A, Table S1), which is somewhat trivial. Food shortage during the developmental period reduces growth and this in turn affects body size later in life [3]. Because body size *per se* has little direct causal effect on fitness in zebra finches [69], more complete developmental canalization for size-related traits may not have evolved. Indeed, despite large variation in mass at day 8 (1.7-12g), the effect of nestling mass on reproductive performance and lifespan was weak, suggesting that fitness is remarkably resilient to variation in early-life conditions [22, 38].

Note that our study is non-experimental and on captive individuals. The latter implies that individuals were kept in a safe environment with *ad libitum* access to food (but with intense social interactions including competition for mates and nest sites). Direct and transgenerational effects on reproductive performance traits may be different in free-living populations, where individuals live and reproduce under potentially more stressful environmental conditions. Additionally, our data set was not ideal to test ‘anticipatory parental effects’. This hypothesis predicts that offspring have higher fitness when the offspring environment matches the parental environment (e.g. [10, 59]). In an ideal experiment, one would manipulate the parents’ and the offspring’s breeding environments in a fully factorial design and examine the effects of matching versus mismatching on offspring performance [8,10,12,59]. Our study only uses observational data and only regarding the similarity of the early growth environments (but not breeding environments). However, a meta-analysis of experimental studies on plants and animals only found a weak trend for small beneficial anticipatory parental effects (effect size d=0.186, highest posterior density: −0.030, 0.393) [59]. Experimental studies are better suited to test causality, but when analyses of observational data suggest no effect, experiments may not provide much insight (note that the 95% confidence interval for the mean effect excluded all biologically relevant effect sizes, e.g. the estimated condition-transfer effects ranged from − 0.017 to 0.002, and anticipatory effects ranged from 0.017 to 0.039). Our approach had the advantage that we could make use of the entire range of observed growth conditions (7-fold difference in mass at day 8), while experimental studies often only induce a 10-15% difference in nestling mass between treatment groups (because ethical concerns prohibit strong treatments; [3, 67]). This then requires larger sample sizes to detect similar phenotypic effects. Our additional confirmatory data sets had smaller sample sizes than the initial data set (Fig 5) and the data were more heterogeneous, because they included individuals from different populations that differ in genetic background, body size and domestication history. It is also noteworthy that the significant effect of father-daughter similarity on daughter tarsus length held up in one confirmatory data set (Fig 5C). Furthermore, one should keep in mind that mass at day 8 is not exclusively a descriptor of the extrinsic growth conditions experienced by the nestling, but may also be influenced by genetic differences among individuals in their ability of convert food into growth. Cross-fostering experiments in the same population indicated a heritability of nestling mass of h^2^ = 0.13 [3].

In summary, for future studies on transgenerational effects, we suggest an approach that renders multiple testing a strength rather than a burden and that consists of four simple steps. (i) Start with clear, one-tailed hypotheses [70], (ii) validation by assessing the direct effects (Fig 1), (iii) meta-analysis of all effects (Fig 2) and – if feasible – (iv) verify the effects with an independent confirmatory dataset (Fig 5). Using this approach, our study shows convincing evidence for small direct effects, and – at best – weak evidence for small transgenerational effects on morphology and fitness. Hence, our study supports the null hypothesis that selection buffers individual fitness against detrimental epigenetic effects, such that the detrimental effects due to stress experienced early in life by the ancestors are not carried on across generations [22, 71].

## Methods

### General procedures

The zebra finch is an abundant, opportunistic breeder in Australia in the wild [72] that also breeds easily in captivity. We used birds from a domesticated zebra finch population with a 13-generation error-free pedigree, maintained at the Max Planck Institute for Ornithology, Seewiesen, Germany (#18 in [73]). We used all individuals (N=2099) for which information was available on laying and hatching dates, egg volume and nestling mass at 8 days of age, both from the focal individual itself, but also from its parents and grandparents. During episodes of breeding in aviaries or in cages, nests were checked daily on weekdays and occasionally during weekends (for further details, see [6,57,74]).

### Early developmental stressors

We examined six parameters that have been used in previous studies as potential proxies of nutrition or stress experienced during early development: (i) the laying order of eggs within a clutch (range: 1-18, mean =3.1, SD=1.7, note that only 5 birds hatched from eggs with laying order >10; such large clutches were found in communal aviary breeding, whereby clutch size was defined as the number of eggs laid consecutively by a focal female allowing for laying gaps of maximally 4 days between subsequent eggs) [53, 54], (ii) the order of clutches laid within a breeding season (range: 1-8, mean =2.1, SD=1.2) [55], (iii) the order of hatching within a brood (range: 1-6, mean =2.1, SD=1.1) [51, 52], (iv) brood size (number of nestlings reaching 8 days of age; range: 1-6, mean =3.3, SD=1.2) [2, 50], (v) relative egg volume (i.e. centered to the mean egg volume laid by a given female; range: −0.26-0.30, mean =0.01, SD=0.07) [56] and (vi) nestling body mass at 8 days of age (range: 1.7-12.4 g, mean =7.2 g, SD=1.6) [3]. Egg volume was calculated as 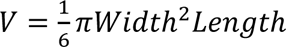, whereby egg length and width were measured to the nearest 0.1mm. Egg volume was not influenced by whether the egg was laid in an aviary or a cage. Nestlings were somewhat heavier in aviaries than in cages (b = 0.19, SE = 0.12, Z = 1.58, P = 0.12).

We hypothesized that individuals (or their parents and grandparents) developed under more stressful conditions if they came from eggs later in the laying, clutch or hatching sequence, were raised in a larger brood, hatched from an egg that was relatively small and had a lower body mass at 8 days of age. For the measure of similarity of parent-offspring early developmental condition (predictor of ‘anticipatory effect’), we calculated the absolute difference in nestling mass at 8 days between parent (mother or father) and offspring (mother-offspring range: 0-8.4 g, mean =1.8 g, SD=1.4; father-offspring range: 0-7.9 g, mean =1.7 g, SD=1.3).

To aid interpretation, we scored all stressors in such a way that all estimated effects are expected to be positive (multiplication by −1 where necessary). Thus, positive effect sizes indicate detrimental effects of a stressor on a trait.

### Morphological and fitness-related traits

We studied the following morphological traits, measured when the individual reached adulthood (median = 115 days of age, range 93-229 days, >95% of birds were 100-137 days old): (i) body mass (measured to the nearest 0.1 g using a digital scale, N = 947 females and 1012 males), (ii) length of the right tarsus (measured from the bent foot to the rear edge of the tarsometatarsus, including the joint, using a wing ruler to the nearest 0.1 mm, N = 944 females and 1008 males; see method 3 in Forstmeier et al. 2007), (iii) length of the flattened right wing (measured with a wing ruler to the nearest 0.5 mm, N = 939 female and 1004 males), (iv) visible clavicular and abdominal fat deposition, scored from 0 to 5 in 0.5 increments (N = 932 females and 989 males), and (v) redness of the beak [57], scored by comparison to a color standard following the Munsell color scale from 0 to 5.5 in 0.1 increments (N = 947 females and 1012 males). Male and female traits were analyzed separately, leading to a total of 10 morphological traits.

We studied the following 13 fitness-related traits (data taken from [6]): (i) female clutch size measured in cages (N = 166 females) or (ii) in aviaries (N = 274 females), (iii) female fecundity, i.e. total number of eggs laid in aviaries without nestling rearing (N = 230 females), (iv) male infertility, measured in cages as the proportion of non-developing eggs (N = 132 males) and (v) in aviaries as the proportion of eggs not fertilized by the social male (N = 237 males), (vi) male siring success, measured in aviaries as the total number of eggs sired (within- and extra-pair; N = 281 males), (vii) female embryo mortality, measured as the proportion of a genetic mother’s embryos dying (N = 228 genetic mothers), (viii) nestling mortality, measured as the proportion of hatchlings in a brood that died before day 35, for a given social mother (N = 233) and (ix) for a given social fathers (N = 228), (x) female and (xi) male seasonal recruits as the total number of independent offspring produced (defined as offspring that survived until day 35; N = 126 males and N =125 females), (xii) female (N = 409) and (xiii) male lifespan (N = 412). In cages, single pairs were kept whereby the partners were assigned to each other; in semi-outdoor aviaries, a group of females and males were kept whereby birds freely formed breeding pairs.

For infertility, embryo and nestling mortality, we used raw data based on the fate of single eggs, while controlling for pseudo-replication by adding male and female identities as random effects in all models (see Statistical Analysis). Female clutch size was analyzed at the clutch level, controlling for female identity, because 94% of females produced multiple clutches. For fecundity, siring success and seasonal recruits, we used the data from individuals within a given breeding season (96% of females and 78% of males had multiple measures for fecundity and siring success, while for seasonal recruits, females and males were only measured once). For easy interpretation of the results, we scored all fitness-related traits in such a way that high trait values refer to better reproductive performance (multiplication by −1 where necessary).

The morphological and fitness-related traits are in general positively correlated within female and male zebra finches (Fig S3 and Table S6).

### Statistics

We estimated the effect of each potential stressor experienced either by the individual itself, or by one of its parents or grandparents on each trait in a separate model (6 stressors x 7 sources x 23 traits = 966 models). We used mixed-effect models and animal models to control for the non-independence of data points due to shared random effects including genetic relatedness. For animal models, we used the package ‘pedigreeMM’ [75] and for mixed-effect models we used ‘lme4’ [76] in R V3.5.1. The 95% CIs of estimated effect sizes were calculated using the ‘glht’ function in the ‘multcomp’ R package while controlling for multiple testing [77], unless stated otherwise.

Morphological traits typically show high heritability, so we included the between-individual relatedness matrix (using pedigree information) as a random effect to control for the genetic relatedness of individuals. In contrast, fitness-related traits typically have low heritability [6], so we analyzed fitness-related traits in mixed-effect models while only controlling for repeated measurements from the same focal individual, parent or grandparent. To compare and summarize the effects of the variables indicating early-life conditions on different traits, we Z-scaled all dependent and all predictor variables (stressors), assuming a Gaussian distribution.

Details on model structures, all scripts and underlying data are provided in the Open Science Framework at https://osf.io/wjg3q/. In brief, for all morphological traits, we fitted sex (male and female), fostering experience (three levels: no cross-fostering, cross-fostered within or between populations), and inbreeding level (pedigree-based inbreeding coefficient, Fped, where outbred birds have Fped=0 and full-sib matings produce birds with Fped=0.25) as fixed effects. For models with beak color, wing and tarsus length as the dependent variable, we also fitted the identity of the observer that measured the trait as a fixed effect to control for between-observer variation. We included the identity of the peer group in which the individual grew up as a random effect. We fitted individual identity twice in the random structure, once linked to the pedigree to control for relatedness between individuals and once to estimate the permanent environmental effect. Additionally, for models with body mass, beak color, wing and tarsus length and fat score as the dependent variable, we included the identity of the batch of birds that were measured together as a random effect (group ID) to control for batch effects between measurement sessions.

For models of fitness-related traits, we controlled for individual age, inbreeding level (Fped), number of days the individual was allowed to breed (in aviaries), the sex ratio (i.e. the proportion of males) and pairing status (force-paired in cages or free-paired in aviaries) by including them as fixed effects, whenever applicable. Additionally, for egg-based models (male fertility, embryo and nestling survival), we controlled for clutch order and laying or hatching order of the egg that was laid/potentially sired by the focal female or male. For models on embryo and nestling survival, we also controlled for the inbreeding level of the offspring. In all models, we included individual identity, breeding season identity, clutch identity, identity of the partner of the focal individual and the pair identity, as appropriate.

We meta-summarized effect sizes using the ‘lm’ function in the R package ‘stats’ V3.6.1, whereby we weighted each effect size by the inverse of the standard error of the estimate to account for the uncertainty of each estimate. Intercepts were removed to estimate the mean of each category unless stated otherwise. First, we meta-summarized the direct effect of each of the six stressors on the individual’s own morphological versus fitness-related traits (‘trait type’, 2 levels). In this model, we fitted the pairwise combination of the trait type and the potential stressor as a fixed effect with 12 levels. Second, we summarized the direct or transgenerational effects (from the individual, its parents and grandparents, 7 levels, ‘stress experienced by a certain individual’) of the most powerful proxy of developmental stress (nestling body mass at 8 days old, see Results) on the morphological versus fitness related traits (2 levels) of males and females (2 levels, ‘sex’). In this model, we fitted the pairwise combination of stress experienced by a certain individual, trait type and sex as a fixed effect with 28 levels. Third, we meta-summarized the transgenerational anticipatory effect of the similarity between parent-offspring in their nestling mass (mother or father in combination with daughters or sons, 4 levels) on the offspring’s morphological versus fitness-related traits (2 levels). Here we included the pairwise combination of parent, offspring sex and trait type as a fixed effect with 8 levels.

Then, we meta-summarized the overall transgenerational effects of conditional transfer and anticipatory effects in two mixed-effect models using the ‘lmer’ function in the R package ‘lme4’ [76], where we weighted each estimate by the multiplicative inverse of its standard error to account for their level of uncertainty. To account for the non-independence between response variables (see Fig S3), we fitted a random effect that reflects their dependencies. For this purpose we grouped all 23 performance traits based on their pairwise correlation coefficients (table S6) into 11 categories (see table S4). The fitted random effect groups the performance traits into 11 categories separately for each ancestor (22 levels for the parents and 44 levels for grandparents). We meta-summarized the overall transgenerational effects of conditional transfer of mass at day 8 experienced by the ancestors (parents and grandparents) on the traits of individuals, by only including an intercept. Last, we meta-summarized the overall transgenerational anticipatory effect of similarity between parent-offspring in their mass at day 8 on the traits of offspring, by only including an intercept.

For visualization, we calculated the expected Z-values with 95% CIs from a normal distribution given the number of Z-values for each group of effects due to each stressor experienced by the focal individual, its mother, its father and its grandparents formulas as follows: expected Z-values as ‘qnorm(ppoints(N Z-values))’ (i.e. the integrated quantiles assuming a uniformly distributed probability of a given number of observations) and 95% CIs of the expected Z-values as ‘qnorm(qbeta(p = (1 ± CI) / 2, shape1 = 1: N Z-values, shape2 = N Z-values:1))’ (i.e. the integrated quantiles of quantiles of a uniformly distributed probability of a given number of observations from a beta distribution) in the R package ‘stats’ V 3.6.1. We visually inspected the ZZ-plots for the expected versus observed Z-values dependent on the direction of the effects. Z-values larger than 1.96 were considered to be significant.

We calculated the sample size for detecting a given effect size with a power of 80% at α=0.05 for a one-tailed hypothesis in the R package ‘pwr’ v1.2-2 [78]. We used the function ‘pwr.r.test’ for observational data and ‘pwr.t.test’ for treatment-based experimental data.

### Confirmatory analysis

For the confirmatory analysis, we used additional birds, including the remaining Seewiesen birds whose maternal nestling mass was known and birds from two other captive populations with short pedigrees: ‘Krakow’ (interbreeding between populations ‘Krakow’ #11 in [73] and ‘Seewiesen’) and ‘Bielefeld’ (wild-derived in the late 1980s, #19 in [73]). To replicate the tests that showed significant effects of maternal early condition and the similarity between mother-daughter early condition on daughter fecundity-related traits (see Results), we used the following samples: (i) female clutch size measured in cages (N = 156 ‘Seewiesen’ and 30 ‘Krakow’ females) or (ii) in aviaries (N = 84 ‘Seewiesen’, 66 ‘Krakow’ and 53 ‘Bielefeld’ females), (iii) female fecundity, measured in aviaries (N = 31 ‘Seewiesen’ females). We Z-scaled nestling body mass at 8 days of age within each population before further analysis, because birds in the recently wild-derived population ‘Bielefeld’ were smaller compared to those of the domesticated ‘Seewiesen’ and ‘Krakow’ populations. We used the ‘lmer’ function from the R package ‘lme4’ to estimate the maternal nestling mass effect on daughters’ fecundity-related traits. The same model structure was used as in the initial tests, but we additionally controlled for between-population differences by including the population where the female came from as a fixed effect. In the model of female fecundity, we removed the variable “number of days the female stayed in the experiment” (because there was no variation) and the random effect “female identity” (because each individual contributed only one data point). To replicate the tests that showed a significant effect of similarity of father-daughter early condition on the daughters’ size-related traits, we used (1) tarsus length of N = 480 ‘Seewiesen’, 290 ‘Krakow’ and 357 ‘Bielefeld’ females and (2) wing length of N = 256 ‘Seewiesen’, 287 ‘Krakow’ and 356 ‘Bielefeld’ females. We analysed the animal model for each population separately, using the same model structure as in the initial test, using the R function ‘pedigreeMM’ from package ‘PedigreemMM’. In the confirmatory models for the Seewiesen population, we removed “author identity” because all birds were measured by the same person.

We meta-summarized the effects of (1) maternal mass at 8 days old, (2) similarity of mother-daughter early condition on her daughters’ fecundity-related traits and (3) similarity of father-daughter early condition on his daughters’ size (see Results) in a ‘lm’ model, by fitting the pairwise combination of test (initial or confirmatory) and the three effects as a fixed effect with 6 levels and the multiplicative reverse of the standard error of each estimate as ‘weight’.

## Data accessibility

Supporting data and R scripts can be found in the Open Science Framework at https://osf.io/wjg3q/ and the fitness-related data can be found at https://osf.io/tgsz8/.

## Authors’ contributions

W.F. and B.K. designed the study. W.F. collected the morphological data. Y.P. and W.F. analyzed the data and interpreted the results with input from B.K. Y.P. and W.F. wrote the manuscript with help from B.K.

## Acknowledgements

We thank M. Schneider for molecular work, and K. Martin, S. Bauer, E. Bodendorfer, J. Didsbury, A. Grötsch, A. Kortner, P. Neubauer, F. Weigel and B. Wörle for animal care and help with breeding.

